# NAIP—NLRC4 Inflammasome Activation in Tuft Cells Activates a PGD_2_–ILC3 Signaling Circuit that Protects Against Enteric Infection

**DOI:** 10.1101/2023.05.11.540443

**Authors:** Madeline J. Churchill, Renate Bauer, Roslyn Honodel, Tighe Christopher, Shuchi Smita, Lindsey Warner, Bridget M. Mooney, Elia D. Tait Wojno, Isabella Rauch

## Abstract

Intestinal epithelial cells (IEC) use innate sensing pathways to distinguish pathogens from commensals. One such pathway, the NAIP—NLRC4 inflammasome, initiates extrusion of infected IEC and mediator release upon cytosolic bacterial sensing. Tuft cells are primarily known for their function in anti-parasite immunity. We previously reported that activation of the inflammasome in tuft cells leads to release of prostaglandin D2 (PGD_2_). We test the hypothesis that tuft cell specific release of PGD_2_ after inflammasome activation initiates antibacterial responses. NAIP—NLRC4 inflammasome activation in tuft cells leads to a type 3 antimicrobial response with increased IL-22 and antimicrobial protein levels within the small intestine, which is dependent on PGD_2_ signaling. A subset of ILC3 express the PGD_2_ receptor CRTH2 and we show them as the source of the increased IL-22. Inflammasome activation in tuft cells also leads to better control of *Salmonella* Typhimurium. These data support that intestinal tuft cells can also induce antibacterial responses.

**Summary:** PGD_2_ release after NAIP—NLRC4 inflammasome activation in tuft cells signals onto ILC3s and mediates host defense mechanisms against *Salmonella* Typhimurium within the small intestine. Tuft cells therefore not only promote immune reactions against parasites, but also bacteria.

## Introduction

The gastrointestinal tract as a barrier tissue is home to many complex interactions between host tissue, the microbiota, and potential pathogenic microbes. One challenge intestinal epithelial cells (IECs) must overcome, is to distinguish commensals from pathogens.

An innate pathway that IECs use to specifically recognize bacterial pathogens is the NAIP— NLRC4 inflammasome. It is activated when NAIP proteins recognize their respective ligands, bacterial T3SS components or flagellin, within the cytosol of the host cell (Kofoed and Vance, 2011; Zhao et al., 2011; Rauch et al., 2016; Zhao et al., 2016; Tenthorey et al., 2014; Rayamajhi et al., 2013; Yang et al., 2013). The strictly cytosolic location of this process ensures distinction between commensal bacteria, which do not invade IEC or use secretion systems to inject effectors, and pathogens.

Induction of the NAIP—NLRC4 inflammasome in intestinal epithelial cells (IEC) is important for host defense against several pathogenic bacteria. By inducing extrusion of the infected epithelial cell, activation of the NAIP—NLRC4 inflammasome in IEC decreases *Salmonella* Typhimurium tissue invasion at acute timepoints, protects against *Citrobacter rodentium*, and renders mice virtually resistant to *Shigella* invasion (Sellin et al., 2014; Rauch et al., 2017; Mitchell et al., 2020; Hausmann et al., 2020; Nordlander et al., 2014; Fattinger et al., 2021). In conjunction with extrusion, inflammasome activation in epithelial cells releases IL-18 and prostaglandin E2 (PGE_2_) a fast-acting lipid inflammatory mediator (Rauch et al., 2017).

These studies determined the impacts of activating the NAIP–NLRC4 inflammasome in all IECs. Inflammasomes are a group of innate sensors activated by different pathogenic insults, that all lead to the same canonical outcome of Caspase-1 activation, cytokine cleavage and pore formation (Churchill et al., 2022). However, it was reported that NLRP6 inflammasome activation in specialized goblet cells causes mucus release (Birchenough et al., 2016), suggesting that inflammasome activation in specific IEC sub-populations can activate different signaling cascades and downstream outcomes.

One rare subset of IEC, called tuft cells, is critical for sensing parasites and signals onto immune cells to initiate anti-parasitic responses within the small intestine (Von Moltke et al., 2016; Nadjsombati et al., 2018; Gerbe et al., 2016; Howitt et al., 2016; McGinty et al., 2020; Lei et al., 2018; Schneider et al., 2018). Tuft cell activation during parasite infection leads to a signaling circuit of tuft cell derived IL-25 and the eicosanoid LTC_4_ to ILC2 (innate lymphoid cells type 2), inducing tuft cell amplification and other extensive remodeling of the intestinal epithelium, required for an efficient anti-parasitic response.

The involvement of tuft cells in the intestinal epithelial sensing of bacterial pathogens and ensuing immune reactions is less understood. Within the lung, release of *P.aeruginosa* quorum sensing molecules activates the bitter sensing taste receptors, Tas2R105 and Tas2R108 on tuft cells to induce Ca^2+^ influx through Trpm5 (Hollenhorst et al., 2022). Tas2R108 also allows for urethral tuft cells to sense heat killed uropathogenic Escheria coli (Deckmann et al., 2014). Within the gastrointestinal tract, tuft cells have been shown to be activated by succinate produced by microbiota (Lei et al., 2018; Banerjee et al., 2020), a ligand which can also be produced by protists and helminths that leads to a type II immune response after tuft cell sensing (Nadjsombati et al., 2018). However, little is known about whether tuft cells specifically respond to pathogenic bacteria. It has been suggested that tuft cells in the intestine sense a specific bacterial ligand, N-C11-G, secreted by the pathogen *Shigella flexneri,* in an invasion independent manner, and this sensing leads to release of eicosanoids such as PGD_2_ (Xiong et al., 2022), a finding that recently has been questioned (Billipp et al., 2023). Synthesis of PGD_2_ occurs when arachidonic acid is metabolized by PTGDS1/2 into PGH_2_, which is further metabolized into PGD_2_ by HPDGS (Funk, 2001). Tuft cells express high levels of eicosanoid synthesis enzymes, including PTGS1 and HPGDS (Sheppe and Edelmann, 2021; Gerbe et al., 2016; Haber et al., 2017; DelGiorno et al., 2020; Bezençon et al., 2008).

Tuft cells also express the NAIP—NLRC4 inflammasome and we have previously shown that activation of this inflammasome specifically in tuft cells mediates the release of PGD_2_ (Oyesola et al., 2021). In the conditions tested, we observe that tuft cells are the only IEC of the small intestine that release PGD_2_ upon inflammasome activation, suggesting an active participation and perhaps tuft cell specific signaling in epithelial bacterial pathogen sensing. During parasitic infection, mice lacking one of the receptors for PGD_2_, CRTH2, expel the helminth parasite *N. brasiliensis* more efficiently than wild type control animals. This observation demonstrates that PGD_2_-CRTH2 has the potential to suppress the anti-helminth response (Oyesola et al., 2021). It is unknown what downstream impacts PGD_2_-CRTH2 signaling might have after NAIP—NLRC4 inflammasome activation during a bacterial infection.

One hint to the function of PGD_2_ during antibacterial responses is that in mice, RNA for the PGD_2_ receptor CRTH2 is expressed by innate lymphoid cells (ILC), ILC3s (Nagashima et al., 2019; Gury-BenAri et al., 2016; Xu et al., 2019). ILC3s are tissue resident cells within the lamina propria that release the cytokines IL-17 and IL-22, which are important for early antibacterial responses, and upregulate antimicrobial protein transcription in IECs (Valeri and Raffatellu, 2016).

Herein, we test the hypothesis that NAIP—NLRC4 inflammasome activation in tuft cells activates a PGD_2_-ILC3 signaling cascade to induce a host antibacterial response. We observe increased small intestinal IL-22 levels after NAIP—NLRC4 inflammasome activation in tuft cells, in a PGD_2_- dependent manner. Activation of this PGD_2_-ILC3 signaling cascade leads to control of *Salmonella* Typhimurium in the small intestine. Inconsistent with (Xiong et al., 2022) we find no changes in CD45+ tuft cells after *Shigella flexneri* infection. Our findings support that tuft cells not only initiate anti-parasitic immunity, but through activation of the NAIP—NLRC4 inflammasome, tuft cells mediate a signaling circuit that activates antibacterial responses.

## Results

To study NAIP—NLRC4 inflammasome activation in tuft cells, we crossed iNLRC4 mice (Rauch et al., 2017) with a tamoxifen inducible tuft cell specific Cre recombinase line (*Pou2f3-Cre^ERT2^*) (Fig. 1A). This crossing yields mice which, upon tamoxifen treatment, express the *Nlrc4* gene only in tuft cells, which we refer to as iNLRC4*-Pou2f3-Cre^ERT2^*.

**Fig. 1.**
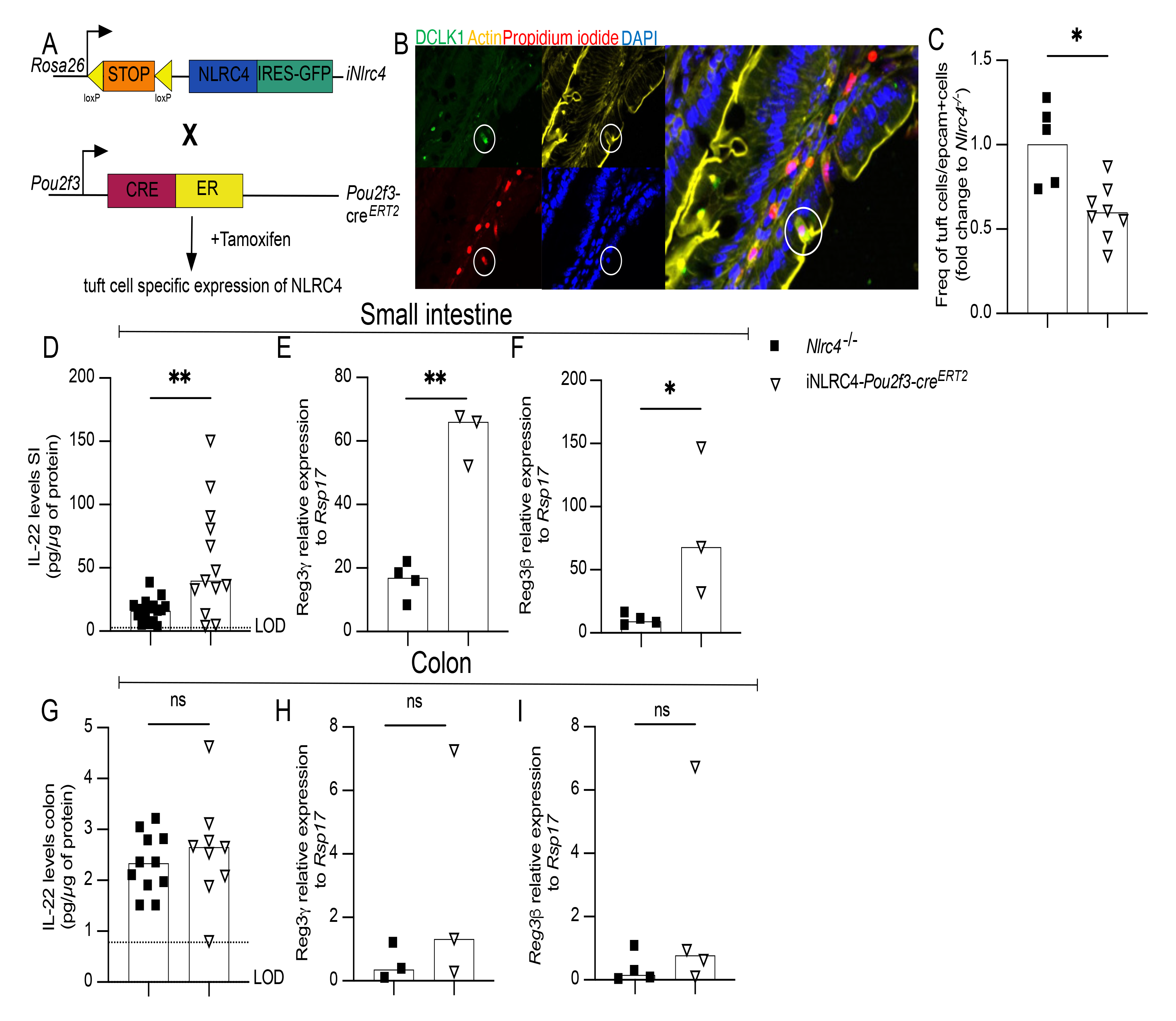
Activation of the NAIP—NLRC4 inflammasome in tuft cells induces an antimicrobial response in small intestinal tissue. (A) Schematic representation of the crossing of iNLRC4 mice with *Pou2f3-cre^ERT2^*mice to yield iNLRC4*-Pou2f3-cre^ERT2^*mice. (B) DCLK-1 (tuft cells: green), Phalloidin (actin: yellow), propidium iodide (pyroptosing cells: red) and DAPI (nuclei: blue) in small intestines from C57BL/6 mice treated with 0.2 µg/g of PA and 0.1 µg/g LFn-Fla for 60 min. (C) iNLRC4*-Pou2f3-Cre^ERT2^* and *Nlrc4^–/–^* littermate controls were retro-orbitally injected with Flatox (LFn-FlaA (0.4ug/g) and PA (0.8ug/g)) and small intestines were harvested 4 hours later. Epithelial cells were isolated from tissues and the number of tuft cells and non-tuft epithelial cells was quantified by flow cytometry. Data was normalized to *Nlrc4^–/–^* mice and is shown as mean (D-I) iNLRC4*-Pou2f3-Cre^ERT2^* and *Nlrc4*^–/–^ littermate controls were retro-orbitally injected with two Flatox dosages (LFn-FlaA (0.4ug/g) and PA (0.8ug/g)) 48 hours apart and tissues were harvested 24 hours after the last injection. IL-22 protein levels within the small intestine (D) and colon (G) were quantified by ELISA, data pooled from multiple experiments. Downstream targets of IL-22 receptor signaling, *Reg3γ* (E, H), and *Reg3β* (F, I) were measured by qPCR and one representative experiment is depicted. Each data point represents one mouse, bars shown as median. Mann-Whitney test was performed to determine significance. All experiments were performed three times. *, P<0.05, **, P<0.001.

We first sought to determine whether tuft cells undergo extrusion after inflammasome activation, as enterocytes do (Sellin et al., 2014; Rauch et al., 2017). Mice were retro-orbitally injected with Flatox (Von Moltke et al., 2016; Rauch et al., 2017), which consists of the NAIP5/6 ligand, flagellin, conjugated to the n-terminal fragment of lethal factor from *B. anthracis*, and protective antigen. Protective antigen forms a pore that allows the flagellin conjugated to the n-terminus of lethal factor to be transported in the cytosol of the host cell and thus selectively activate the NAIP— NLRC4 inflammasome. We observe tuft cells (DCLK1+) positive for propidium iodide, signifying pore formation after inflammasome activation, and purse-like remodeling of actin demonstrating extrusion around tuft cells in intestines of wildtype mice injected with FlaTox (Fig. 1B). Comparing iNLRC4*-Pou2f3-Cre^ERT2^* mice to inflammasome deficient mice, the percentage of tuft cells detectable by flow cytometry among IEC is significantly decreased 4 hours after inflammasome activation specifically in tuft cells (Fig. 1C). Taken together these data support that tuft cells are extruded from the epithelial layer after tuft cell activation of the NAIP—NLRC4 inflammasome.

We have previously shown that PGD_2_ is released after NAIP—NLRC4 inflammasome activation in tuft cells (Oyesola et al., 2021). Since lipid mediators like PGD_2,_ are released quickly after activation, we hypothesized that PGD_2_ release would occur proceeding or during cell extrusion, allowing signaling to nearby cells. Small intestinal ILC3 from the lamina propria express CRTH2 by RNA-seq analysis (Nagashima et al., 2019; Xu et al., 2019). Therefore, we hypothesized that tuft cell release of PGD_2_ would signal to activate ILC3 during acute bacterial infection. To study a potential tuft cell-PGD_2_-ILC3 communication axis, we first analyzed levels of one of the ILC3 signature cytokines, IL-22, in intestinal tissue after tuft cell inflammasome activation. After two doses of Flatox spaced 48 hours apart, we observe a significant increase in IL-22 protein levels within the small intestine (duodenum, jejunum and ileum) in iNLRC4*-Pou2f3-Cre^ERT2^*mice compared to *Nlrc4^–/–^* littermates (Fig.1D). In conjunction, transcription of antimicrobial proteins downstream of IL-22 signaling, *Reg3γ* (Fig. 1E), and *Reg3β* (Fig. 1F) is significantly increased in iNLRC4*-Pou2f3-Cre^ERT2^* mice compared to *Nlrc4^–/–^* littermates after inflammasome activation. IL- 22 protein levels, *Reg3γ*, and *Reg3β* were not significantly different in the colon (Fig. 1G-1I), consistent with previous literature that demonstrates that small intestinal and colonic tuft cells can have different functions (Strine and Wilen, 2022). These results demonstrate that IL-22, an ILC3 signature cytokine, and antimicrobial transcripts within the small intestine are increased after NAIP—NLRC4 inflammasome activation in tuft cells.

To determine if ILC3s are the source of IL-22 within the small intestine after tuft cell inflammasome activation, we used flow cytometry to measure intracellular IL-22 levels in various cell types that have been shown to produce IL-22 (Perusina Lanfranca et al., 2016). We identified the subtypes of ILC populations using a previously developed flow cytometry panel (Pokrovskii et al., 2019; Talbot et al., 2020) (Supplemental 1A, Fig. 2A). There was a significant increase of IL-22 expressing ILC3s (Fig. 2A-C) in iNLRC4*-Pou2f3-Cre^ERT2^* mice compared to *Nlrc4^–/–^*littermates after inflammasome activation. We observe no significant differences in the number of IL-22+ TCR*β*+ or TCR*γ/δ*+ T cells (Fig. 2D, E). These data suggest that ILC3s are the cell type responsible for increased IL-22 after tuft cell specific inflammasome activation.

**Fig. 2.**
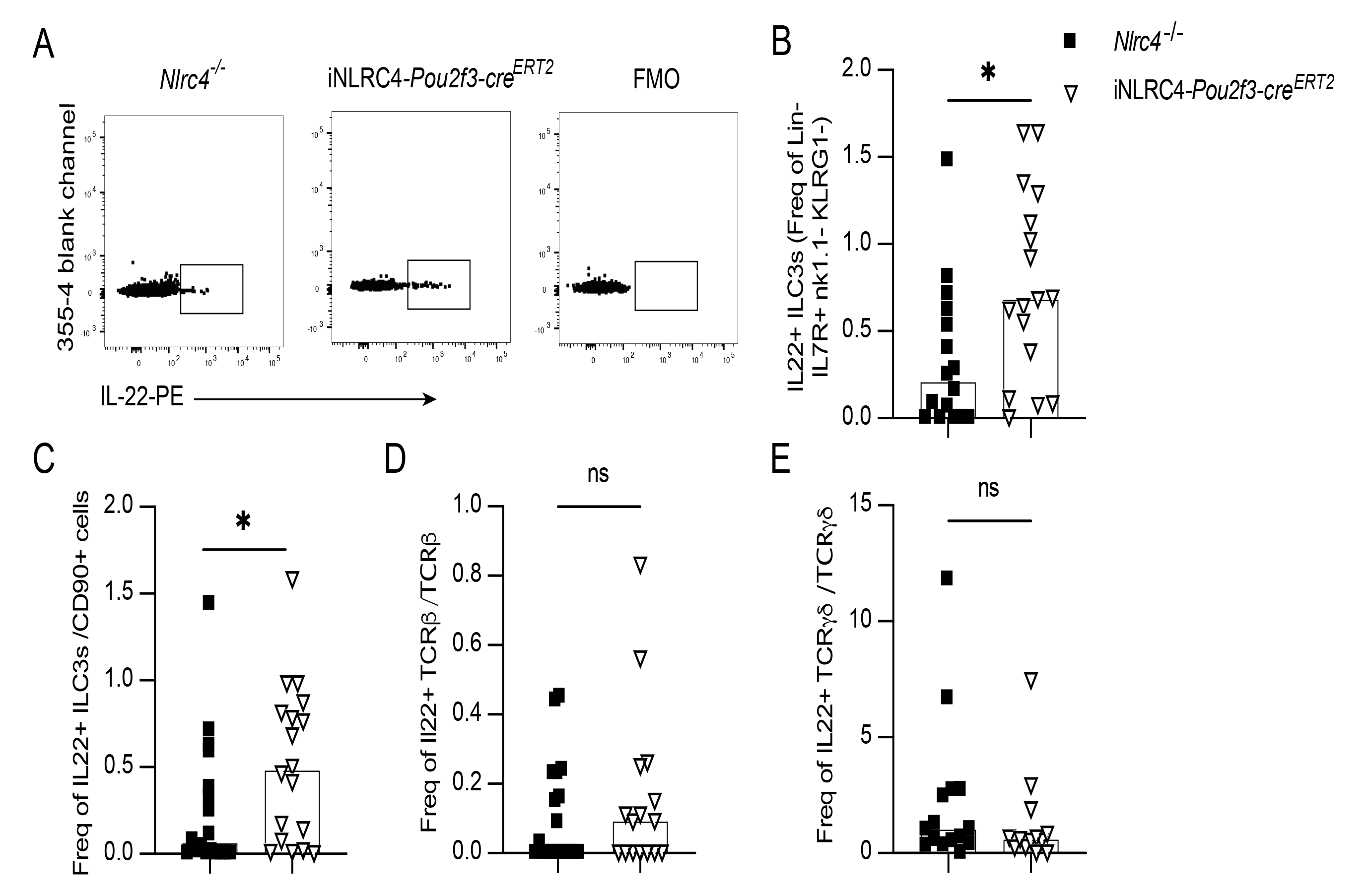
ILC3s are the source of increased IL-22 within the small intestine after NAIP—NLRC4 inflammasome activation in tuft cells. iNLRC4*-Pou2f3-cre^ERT2^* mice or littermate controls were administered two dosages Flatox (LFn-FlaA (0.4ug/g) and PA (0.8ug/g)) retro-orbitally 48 hours apart and small intestine was harvested 24 hours after last Flatox injection. Cells from the lamina propria were isolated and intracellular IL-22 was measured by flow cytometry. (A) Representative flow plots of intracellular IL-22 in ILC3s from *Nlrc4^–/–^*, iNLRC4*-Pou2f3-cre^ERT2^* and FMO controls. (B) Quantification of IL-22+ ILC3s as in (A). Frequency of IL-22+ ILC3s among CD90+ cells (C). Frequency of IL-22+ TCR*β*^+^ cells (D). Frequency of IL-22+ TCR*γ/δ* ^+^ cells (E). Each data point represents one mouse, pooled data from multiple experiments, and bars shown as median. Mann-Whitney U test was performed. All experiments were performed three times. *, P<0.05.

To confirm the observation of PGD_2_ receptor expression on ILC3s by single cell RNA sequencing we determined if the *Gpr44* locus, which encodes the CRTH2 protein, is active in ILC3s. We used a novel *Gpr44* transcriptional reporter mouse. These mice have a floxed coding sequence of *Gpr44* with proximal GFP, which will be expressed once the *Gpr44* locus is deleted (Supplemental Fig. 2.). Using these mice crossed to *Rorγt-Cre* mice, we observe a percentage of CCR6 negative ILC3s with active *Gpr44* expression within the small intestine during steady state (Fig. 3A, B). To determine if the observed IL-22 increase after tuft cell inflammasome activation within the small intestine is dependent on CRTH2 signaling, we administered an inhibitor specific to this receptor, AZD1981. Treatment with AZD1981 completely abrogated the increase in small intestine IL-22 in iNLRC4*-Pou2f3-Cre^ERT2^* mice after tuft cell inflammasome activation (Fig. 3C). AZD1981 also blocks increases in *Reg3γ* (Fig. 3D) and *Reg3β* (Fig. 3E) within the small intestine of iNLRC4*- Pou2f3-Cre^ERT2^* mice. These data suggest that the increases in IL-22 and its downstream targets after tuft cell inflammasome activation are dependent on PGD_2_ signaling on ILC3.

**Fig. 3.**
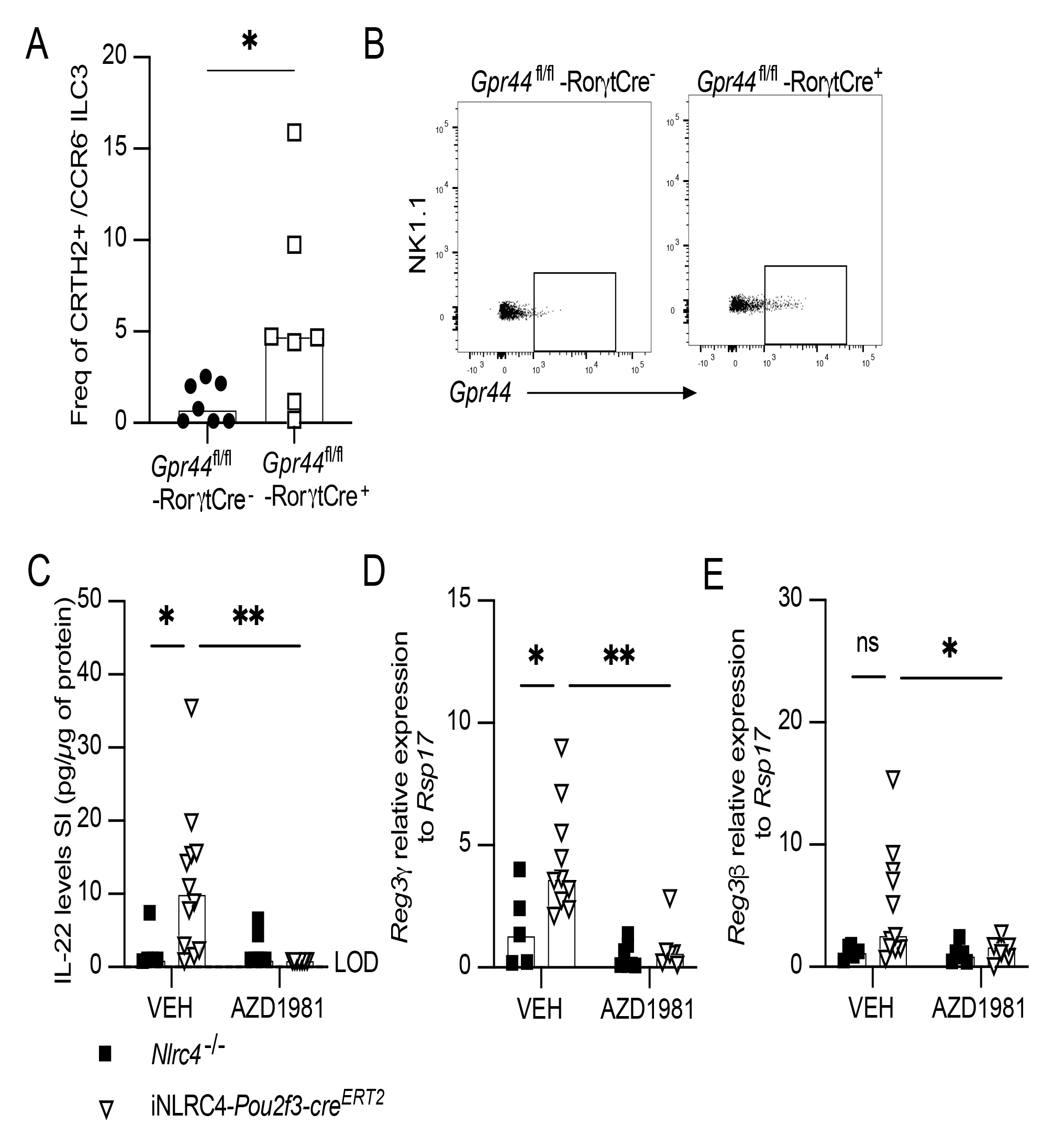
The antibacterial signature in the small intestine after tuft cell NAIP—NLRC4 inflammasome activation is dependent on CRTH2 signaling. (A) Lamina propria cells from *Gpr44*-GFP mice were analyzed by flow cytometry and the percentage of GFP+ CCR6- ILC3s determined. Significance was determined by Mann-Whitney U Test. (B) Representative flow cytometry plots of GFP+ ILC3s from (A). iNLRC4*-Pou2f3-cre^ERT2^*mice and *Nlrc4^–/–^*littermate controls were retro-orbital injected with two dosages of Flatox (LFn-FlaA (0.4ug/g) and PA (0.8ug/g)) 48 hours apart, 30 minutes after intraperitoneal injection of AZD1981 or vehicle. The tissue was harvested 24 hours after the final Flatox injection. (C) IL-22 protein levels were measured in the small intestine by ELISA. *Reg3γ* (D)and *Reg3β* (E) were measured by qPCR in the small intestine. Each data point represents one mouse, pooled data from multiple experiments, bars shown as median. Two-way ANOVA corrected for multiple comparisons in C, D and E. All experiments were performed at least twice. *, P<0.05, **, P<0.01, ***P<0.001.

How intestinal tuft cells contribute to host defense responses during bacterial infection remains incompletely understood. A previous report suggested that tuft cells can sense a secreted extracellular bacterial ligand (NOT invasion) and trigger a feedback cycle leading to tuft cell proliferation during infection with *Shigella flexneri* (Xiong et al., 2022). In our hands, we did not observe an increase in total tuft cells (Supplemental Fig. 3A, B) or CD45+ tuft cells after *Shigella flexneri* infection (Suppl. Fig. 3C, D). To determine if tuft cell inflammasome signaling (which senses active invasion by a bacterium) controls pathogenic bacteria, we infected iNLRC4*-Pou2f3-Cre^ERT2^* mice and littermate *Nlrc4*^–/–^ controls with *Salmonella* Typhimurium ΔssaR. These bacteria lack a component of the SPI-2 T3SS, and therefore cannot successfully evade the phagosome of myeloid cells (Pfeifer et al., 1999), which leads to productive infection of only epithelial cells via the SPI-I T3SS. This allows us to study a model gastrointestinal pathogen without the complications of systemic spread and mortality at later timepoints (Matheis et al., 2020; Gül et al., 2023). At 18 hours or 5 days post infection, the ileum and cecum were harvested. Tissue colony forming units (CFUs) were determined from gentamicin treated and washed samples to eliminate luminal bacteria. We hypothesized that, as tuft cells are rare in SPF mice, their extrusion will not significantly affect pathogen numbers, however PGD_2_ signaling induced antimicrobial signaling could influence pathogen clearance at later timepoints. Indeed, there is no significant difference in CFUs in either the ileum (Fig. 4A) or cecum (Fig. 4B) at 18 hours post infection, a timepoint where enterocyte inflammasome induced extrusion shows the strongest effect in invasion prevention (Sellin et al., 2014; Fattinger et al., 2021; Rauch et al., 2017). A trend for a decrease in CFU is, however, appreciable in the small intestine (Fig. 4A). At 5 days post infection, iNLRC4*- Pou2f3-Cre^ERT2^* mice have a significantly decreased CFU count compared to *Nlrc4^–/–^* within the small intestine (Fig. 4C). No differences were observed in the cecum (Fig. 4D). 50% of iNLRC4*- Pou2f3-Cre^ERT2^* animals completely cleared the bacteria from their small intestine at day 5 post infection, compared to only 25% of *Nlrc4^–/–^* mice (Fig. 4E). Consistent with our earlier result with FlaTox inflammasome activation, we observed increased IL-22 protein levels within the small intestine of iNLRC4*-Pou2f3-Cre^ERT2^* mice compared to *Nlrc4^–/–^*littermate controls at 5 days post infection (Fig. 4F). No differences in IL-22 were observed in the cecum (Fig. 4G). Based on these data, we conclude that PGD_2_-ILC3 signaling after tuft cell specific inflammasome activation can contribute to bacterial pathogen clearance in the small intestine.

**Fig. 4.**
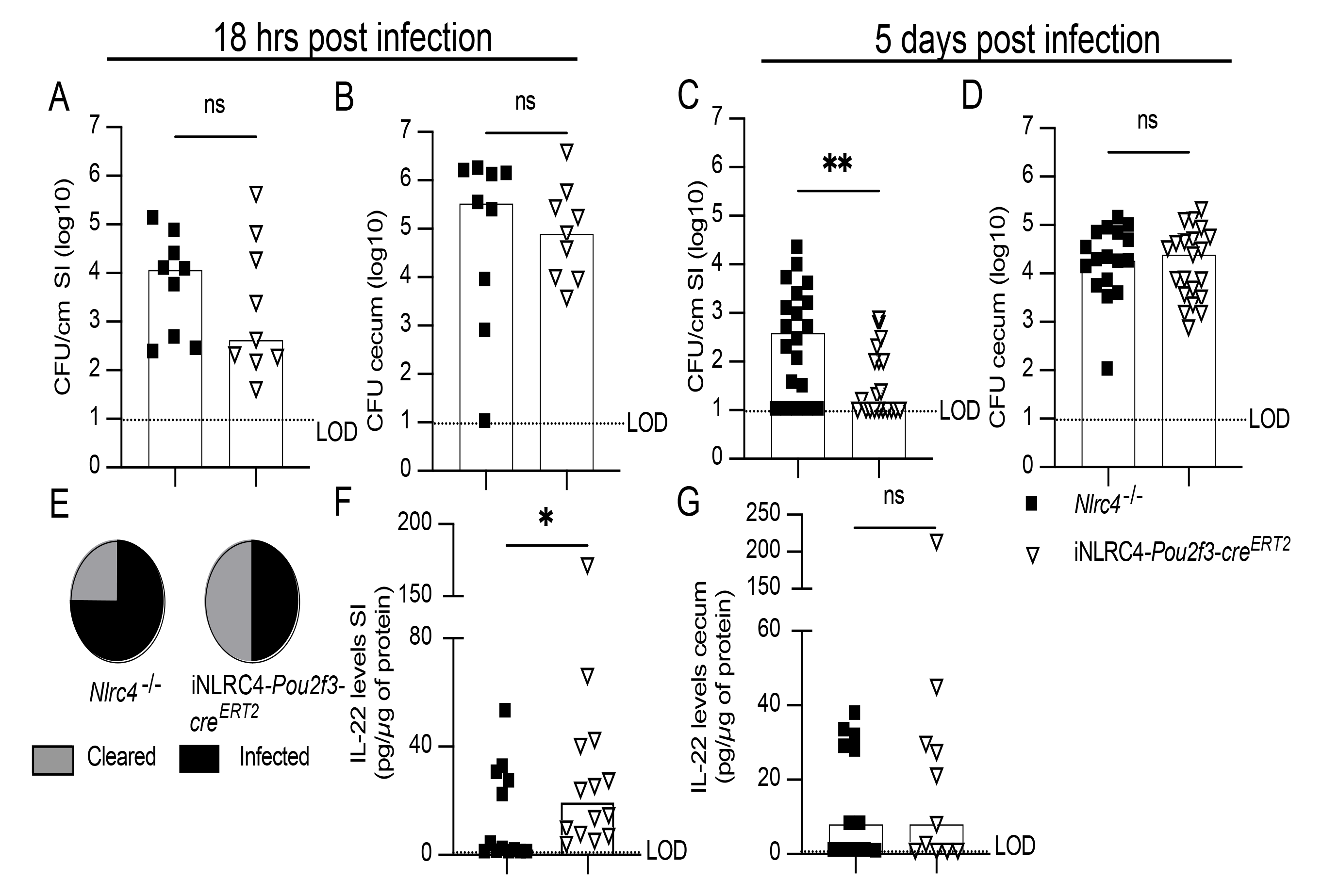
Tuft cell specific NAIP—NLRC4 inflammasome activation mediates control of *Salmonella* Typhimurium infection in the small intestine. iNLRC4*-Pou2f3-cre^ERT2^ mice* or *Nlrc4*^–/–^ littermate controls were pre-treated with oral gavage of streptomycin, then 24 hours later infected with 5×10^7^ CFUs of *Salmonella* Typhimurium ssrA by oral gavage. Tissue CFUs from small intestine (A) and cecum (B) 18 hours post infection and small intestine (C) and cecum (D) 5 days post infection. (E) Number of *Nlrc4^–/–^* or iNLRC4*-Pou2f3-cre^ERT2^* that had no detectable CFUs at day 5 post infection. IL-22 protein levels were determined by ELISA in small intestine (F) or cecum (G). Each data point represents one mouse, pooled data from multiple experiments, bars shown as median. Significance was determined by Mann-Whitney U test (A-D, F-G). All experiments were performed at least twice *, P<0.05, **, P<0.01.

## Discussion

The functions of intestinal tuft cells are not completely known. Understanding how different cell types, including tuft cells within the gastrointestinal tract contribute to the host inflammatory response will provide more foundational knowledge to understand how inflammatory responses are initiated, maintained or changed within the gastrointestinal tract. It may also uncover new potential therapies for enteric disease.

We demonstrate that NAIP—NLRC4 inflammasome activation in tuft cells induces a PGD_2_ – ILC3 signaling pathway within the small intestine that promotes clearance of a gastrointestinal bacterial pathogen. We demonstrate that tuft cells are not solely parasitic sensors, but also participate in host defense responses against bacteria. We have previously shown that tuft cells release PGD_2_ after NAIP—NLRC4 inflammasome activation (Oyesola et al., 2021). While PGD_2_ is detrimental to antiparasitic responses, our data supports that PGD_2_ release during bacterial infection is beneficial to the host and leads to better control of the pathogenic bacteria. It is tempting to speculate that PGD_2_ signaling in the intestine leads to a balance between anti-parasitic responses and anti-bacterial responses, and that tuft cells are the orchestrators of this balance. Tuft cells are also preferentially infected by murine norovirus (Wilen et al., 2018). It is conceivable that tuft cells integrate many different pathogen signals and have the potential to act as a rheostat for the ensuing host inflammatory response.

A previous report demonstrated tuft cell proliferation and PGD_2_ release upon sensing of a bacterial metabolite, N-C11-G, during *Shigella flexneri* infection, and this promoted tuft cell amplification and protection (Xiong et al., 2022). While we also observe that small intestinal tuft cells release PGD_2_ (Oyesola et al., 2021), the *Shigella* study did not rule out the involvement of the NAIP— NLRC4 inflammasome during infection with NAIP ligand expressing pathogens as we have shown here. In addition, in our hands we do not observe an increase in CD45+ tuft cells during *Shigella* infection. This is consistent with previous observations that *Shigella* cannot efficiently colonize inflammasome competent mice due to extremely efficient restriction of these bacteria by the acute epithelial extrusion response (Mitchell et al., 2020). In agreement with our observation, a recent study reported that administration of N-C11-G to intestinal epithelial monolayers did not result in release of leukotrienes or cause ion flux indicating that tuft cells did not respond to this ligand (Billipp et al., 2023). Thus, the recognition of pathogenic bacteria specific extracellular ligands by tuft cells warrants further study.

Here we link inflammasome triggered tuft cell PGD_2_ release to activation of ILC3s. The crosstalk between lipid mediator release from tuft cells and innate lymphoid cells is reminiscent of signaling observed during anti-parasitic responses. During *H. polygyrus* infection, a type of leukotriene, LTC_4_, is released by tuft cells, and induces ILC2s to proliferate and release IL-13 (McGinty et al., 2020). Using more sensitive methods than RNA-sequencing, *Gpr44* expression is also detectable on goblet cells and tuft cells (Oyesola et al., 2021). Therefore, additional effects of PGD_2_ released by tuft cells, especially during type 2 inflammation, where goblet and tuft cells are increased, are conceivable. Our findings suggest that tuft cells have a unique ability to sense multiple different pathogens and by release of specific lipid mediators, can manipulate downstream inflammatory responses.

## Methods

### Mice, Flatox injections and inhibitor treatment

iNLRC4 (Rauch et al., 2017), *Nlrc4^–/–^* (Tenthorey et al., 2020), *Pou2f3-Cre^ERT2^* (McGinty et al., 2020), CRTH2^fl/fl^*-*GFP mice, B6.FVB-Tg(Rorc-cre)1Litt/J (common name: *Rorγt-Cre* Strain # 022791 (Eberl et al., 2004)) breeding was maintained in specific-pathogen-free facilities at OHSU. iNLRC4*-Pou2f3-cre^ERT2^* mice were generated by crossing iNLRC4 mice with *Pou2f3-Cre^ERT2^*, both on a *Nlrc4^–/–^* background. To determine if the *Gpr44* locus is active in ILC3s, CRTH2^fl/fl^ mice were crossed with *Rorγt-Cre* mice. To induce Cre^ERT2^, iNLRC4*-Pou2f3cre^ERT2^*and *Nlrc4^–/–^*littermate mice were given ad libitum access to tamoxifen impregnated food (Envigo, 500 mg/kg, TD.130857) for one week before experiment began and remained for the duration of the experimentation. One week after tamoxifen food administration, mice were retro-orbitally injected with Flatox comprised of LFn-FlaA from *V.parahemolyticus* (0.4ug/g) and PA (0.8ug/g) given at 4ul/g of body weight.

AZD1981 or Vehicle (1:5 DMSO:PBS) was given 5ug/g body weight. For all *in vivo* studies, age-and sex-matched, cohoused 8-to-20-week littermate or F2 (Robertson et al., 2019) mice were used and all experiments repeated at least twice.

For detection of pyroptotic epithelial cells, mice were retro-orbitally injected with 100µg propidium iodide 10 minutes before euthanasia.

### Generation and validation of CRTH2 F/F-GFP mice

The exon 3 coding region of the *Gpr44* gene (NCBI Reference Sequence: NM_009962.3) on mouse chromosome 19 was targeted for conditional deletion. To construct the targeting vector, mouse genomic fragments containing homology arms were amplified from a BAC clone using high fidelity Taq DNA polymerase and were sequentially assembled into a targeting vector with the polyA-loxP-endogenous SA of intron 2-5’UTR of exon 3-EGFP-SV40 polyA cassette. In the targeting vector, the Neo cassette was flanked by SDA (self-deletion anchor) sites and DTA was used for negative selection. The targeting construct was linearized by restriction digestion, purified, and transfected into C57BL/6 ES cells according to Taconic Inc. (Petersburg, NY) proprietary methods. The transfected ES cells were selected with G418 (200 μg/mL). G418 resistant clones (188) were picked, amplified, and PCR screened for homologous recombination, identifying 11 potential targeted clones. Six were further confirmed by Southern blot analysis to be correctly targeted. Targeted ES cell clones were injected into C57BL/6 albino embryos, which were then re-implanted into CD-1 pseudo-pregnant females. Founder animals were identified by their coat color and germline transmission was confirmed by breeding with C57BL/6 females and subsequent genotyping of the offspring.

CRTH2 F/F-GFP mice were then bred at the University of Washington to Vil1-Cre mice to generate Vil1-Cre x CRTH2 F/F-GFP animals. To test cell lineage-specific deletion of *Gpr44* in Vil1-expressing intestinal epithelial cells, small intestinal epithelial cells and splenocytes were isolated as described previously (Oyesola et al., 2021) from Vil1-Cre+ CRTH2 F/F-GFP, Vil1-Cre+ CRTH2 WT/WT, and Vil1-Cre-CRTH2 F/F-GFP mice. DNA was extracted using the GenCatchPCR Purification kit (Epoch Life Science, Misouri City, TX) according to manufacturer instructions. A genotyping PCR was used to amplify genomic DNA from the WT allele (239 bp); the targeted, non-recombined allele (348 bp); and the targeted, recombined allele (279 bp) to confirm recombination only in Vil1-Cre CRTH2 F/F-GFP mice in small intestine IECs (Sup. Fig. 2). Primers used were *F1: 5’-GTTGGATCCACTGGAACTGAATTAT-3’* and *R1: 5’- TGTAATCTCAGTGCTGGAGA AAGAC-3’*.

All animal procedures were approved and performed in accordance with the Institutional Animal Care and Use Committees at OHSU and the University of Washington.

### Bacterial Infections

The day before infection, mice were food restricted for 4 hours, then orally gavaged with 100μL of 250mg/ml streptomycin sulfate in H_2_O to deplete the microbiome (Barthel et al., 2003). On the day of infection, mice were again food restricted for 4 hours and then infected by oral gavage with either 5×10^7^ CFUs of *Salmonella* Typhimurium ssrA in 100μL PBS or 5×10^6^ CFUs of *Shigella flexneri* in 100μL PBS. Ileum and ceca were collected from *Salmonella* Typhimurium ssrA infected mice 18 hours or 5 days post infection, and processed for CFU enumeration as described below or protein analysis. Whole small intestines from *Shigella flexneri* mice were harvested at 2 days post infection and processed for flow cytometry analysis.

### FlaTox component production

LFn-FlaA. *V. parahaemolyticus* was produced in insect cells as previously described (Mitchell et al., 2020). Briefly, using the BaculoGold system, recombinant protein for cytosolic delivery of *Vibrio parahaemolyticus* FlaA was produced. The FlaA coding sequence was subcloned into pAcSG2-6xHis-LFn and then co-transfected with the BestBac linearized baculovirus DNA (Expression systems) into SF9 cells. Infectious baculovirus was then incubated with High Five cells to generate recombinant proteins. Cells were lysed on ice using a dounce homogenizer in lysis buffer. Supernatants from clarified samples were bound to nickel resin, then recombinant proteins were column purified by gravity.

Protective antigen (PA) was purified as described before (von Moltke et al., 2012). Briefly, PA was expressed in *E.coli* periplasm and purified using periplasmic lysis, followed by dialysis and anion exchange chromatography. Endotoxin was removed using Pierce endotoxin removal columns.

### Harvesting tissue

Upon completion of the experimental timeline, mice were euthanized by CO_2_ asphyxiation and cervical dislocation was performed to ensure euthanasia. An incision was made in the peritoneum of mice and the small intestine was cut from the cecum. For flow cytometry, the whole small intestine was collected, fileted open, rinsed in PBS and cut into 2 cm pieces. For protein and RNA quantification, the whole small intestine was removed and then cut into three 3 pieces (separating the duodenum, jejunum, and ileum). Feces were removed and 2 representative pieces from each region were flash frozen and processed for either protein or qPCR analysis. Another 2 pieces from each region were fileted open and pinned flat into shallow dishes containing 4% paraformaldehyde for 2 hrs at RT. The fixed tissue was placed into 30% sucrose overnight, then rolled into a swiss roll and placed into cassettes containing OCT and frozen until ready for cutting.

### CFU enumeration

Briefly, 2 cm pieces of distal small intestine and half of the cecum was collected from *Salmonella* Typhimurium infected mice and washed in 400μg/ml gentamicin in PBS for 30 minutes at room temperature. To ensure that gentamicin was sufficiently washed away, tissue was rinsed with ice-cold PBS five times. Tissue was then homogenized in ice-cold PBS with a Kinematica Polytron. Serial dilutions were made of cecum and 4 x 10 μL droplets from each serial dilution were plated on MacConkey agar plates containing streptomycin. 100μL of undiluted small intestine homogenate were plated directly onto MacConkey agar using bead plating. 18-20 hours later, colonies were enumerated and final CFUs was determine for each tissue type.

### Cryostat and IF staining

OCT embedded sections were cut at 7μm and placed onto charged glass slides. The tissue was placed into PBS+0.5% Tween for 3 minutes. Tissue was rinsed and then tissue sections were incubated with PBS+0.1% Triton with 10% goat serum for 30 minutes at room temperature. Sections were then immediately incubated with primary antibodies (DCLK-1, ABcam ab31704 rabbit anti-mouse 1:1000 in PBS+0.5% Tween) overnight at 4 degrees C. Samples were rinsed with PBS+0.5% Tween 3 × 5 minutes. Secondary antibodies were incubated for 1 hour at room temperature (goat anti-rabbit 488 Jackson ImmunoResearch Labs, 1:500 in PBS+0.5% Tween). Samples were rinsed with PBS+0.5% Tween 3 × 5 minutes. Phalloidin-647 (Thermo Fisher, A22287, 0.165μM) was incubated on tissue for 30 minutes at RT. Then samples were incubated in DAPI (0.1μg/ml in PBS) for 10 minutes at room temperature, then cover-slipped. Samples were imaged using the Zeiss Apotome.

### Protein Analysis

Flash frozen tissue was placed into PBS+ cOmplete mini protease inhibitor cocktail (Milipore-Sigma #4693124001) and homogenized with a handheld Kinematica Polytron. Samples were frozen and then thawed fully before they were spun down at 10,000 x g to pellet cellular debris. Protein concentration of the supernatant was determined by following manufacturers instruction for the Pierce BCA Protein Assay Kit (ThermoFisher-Scientific #23227). Once total protein concentration was determined, 10 μgs of protein were loaded into manually coated IL-22 96-well High Affinity plates and IL-22 levels were determined by IL-22 ELISA kit (ThermoFisher-Scientific #88-7422-22) following manufacturer’s protocol. The 96-well plate was read using a plate reader (CLARIOstar).

### qPCR analysis

RNA from tissue was purified using the TRIZOL method by manufacturer’s protocol (ThermoFisher Scientific). Samples were measured for RNA concentration and purity. *cDNA*: 1 μg of RNA was incubated with 1μL of DNAase in 10μL total and allowed to incubate for 30 minutes at 37 C. DNAse stop solution was added and incubated for 10 minutes at 65 C. dNTPs were incubated for 5 minutes at 65 C and then cooled to 4 C. Master mix containing Superscript IV reverse transcriptase (ThermoFisher Scientific #18090050) was added to samples and incubated at 55 C for 10 minutes, followed by 80 C for 10 minutes. *qPCR analysis:* Real-Time PCR on cDNA was performed using PowerUp SYBR green Master mix (ThermoFisher Scientific #A25742) and analyzed with an Quantstudio3 (Applied Biosystems). Relative quantities were determined by normalization to housekeeping gene.

Primer sequences: *Reg3γ* F: TTCCTGTCCTCCATGATCAAAA R: CATCCACCTCTGTTGGGTTCA, *Reg3β F:* ATGCTGCTCTCCTGCCTGATG R: CTAATGCGTGCGGAGGGTATATTC, Rps17 (housekeeping gene) F: CGC CAT TAT CCC CAG CAA G R: TGT CGG GAT CCA CCT CAA TG

### Flow Cytometry

#### For Epithelial cell isolations

Isolated small intestines were placed into 15 ml of DTT wash solution (5% FCS, 10mM HEPES, 5mM DTT, HBSS (no Ca/Mg)) gently shaking at 37 C for 10 minutes. Samples were vortexed and then placed into EDTA wash solution (5% FCS, 10mM HEPES, 5mM EDTA, HBSS (no Ca/Mg)) gently shaking for 15 minutes at 37 C. Supernatant was collected through a 70 μm cell strainer. Another wash of EDTA wash solution was added to small intestines and incubated for 5 minutes while gently shaking at 37 C. The second epithelial cell isolation was collected and added to the previous epithelial cell flow through. Samples were then stained for: Live dead (Zombie Aqua), Epcam -AF647, CD24-PE, C-kit-PerCp, CD45+-AF700 and SiglecF-BV421. Tuft cells were identified as Epcam+, SiglecF+ and CD24+ cells.

#### For Lamina Propria Isolations

Lamina propria single cell suspensions were collected using the lamina propria isolation protocol from Miltenyi Biotech with modifications. Briefly, isolated small intestines were washed in ice cold PBS, and then placed into strip buffer (5% FBS, 10mM HEPES, 5mM DTT and 5mM EDTA in PBS) gently shaking at 37 C for 20 minutes. Samples were vortexed and then placed into a second wash with strip buffer gently shaking for 20 minutes. Samples were rinsed with D10 media (10% FBS, 10mM HEPES in DMEM), then digested with the Lamina Propria Isolation kit from Miltenyi Biotech following manufacturers protocol while gently shaking for 30 minutes. Samples were then placed into C tubes and mechanically digested using the Miltenyi Biotech tissue homogenizer – small intestine program. The digested samples were then put through a 70μm cell strainer and 4×10^6^ cells stained with: Intracellular IL-22 Flow panel: Zombie Blue for live dead stain, NK1.1-APC CD45-AF700, CD19-BV650, CD14-BV650, CD11c-BV650, CD3e-BUV395, CD90- BUV661, Rorgt-BUV711, IL7Ra-BV421, TCRa/b-BV510, TCRy/d-FITC, and IL-22-PE. ILC3s were identified as lin-CD90+IL7Ra+Nk1.1-KLRG1-. Samples were run on BD Symphony and analyzed using FlowJo (BD, Ashland OR). ILC3 staining for CRTH2 expression: Zombie Blue, CRTH2-GFP, NK1.1-APC CD45-AF700, CD19-BV650, CD14-BV650, CD11c-BV650, CD3e-BUV395, CD90-BUV661, IL7Ra-BV421, TCRα/β-BV510, TCR*γ*/*δ*- BV510, CCR6-BV605.

## Acknowledgments

The authors thank members of the Rauch laboratory and Jakob von Moltke for their input and Patrick Mitchell (University of Washington, Seattle, Washington) and Leigh Knodler (The University of Vermont) for reagents. The authors further wish to thanks members of the OHSU flow cytometry and microscopy cores for support and advice.

Author contributions: M.J. Churchill and I. Rauch designed, performed, and analyzed experiments. R. Bauer, R. Honodel, B.M. Mooney, L. Warner, S. Smita, T. Christopher and E.D. Tait Wojno performed, analyzed, and/or assisted with experiments and contributed data. M.J. Churchill, I. Rauch and E.D. Tait Wojno made significant intellectual contributions. M.J. Churchill and I.Rauch wrote the manuscript with input from all coauthors. I.Rauch supervised the project.

This work was supported by the National Institutes of Health, National Institute of Allergy and Infectious Diseases (R01 AI167974) and the Rainin Foundation Synergy Award to I. Rauch; the National Institutes of Health (R01 AI130379 and R01 AI167974) to E.D. Tait Wojno, Oregon Health and Science University funds to I. Rauch; and University of Washington startup funds to E.D. Tait Wojno.

---

**Supplemental Fig. 1.**
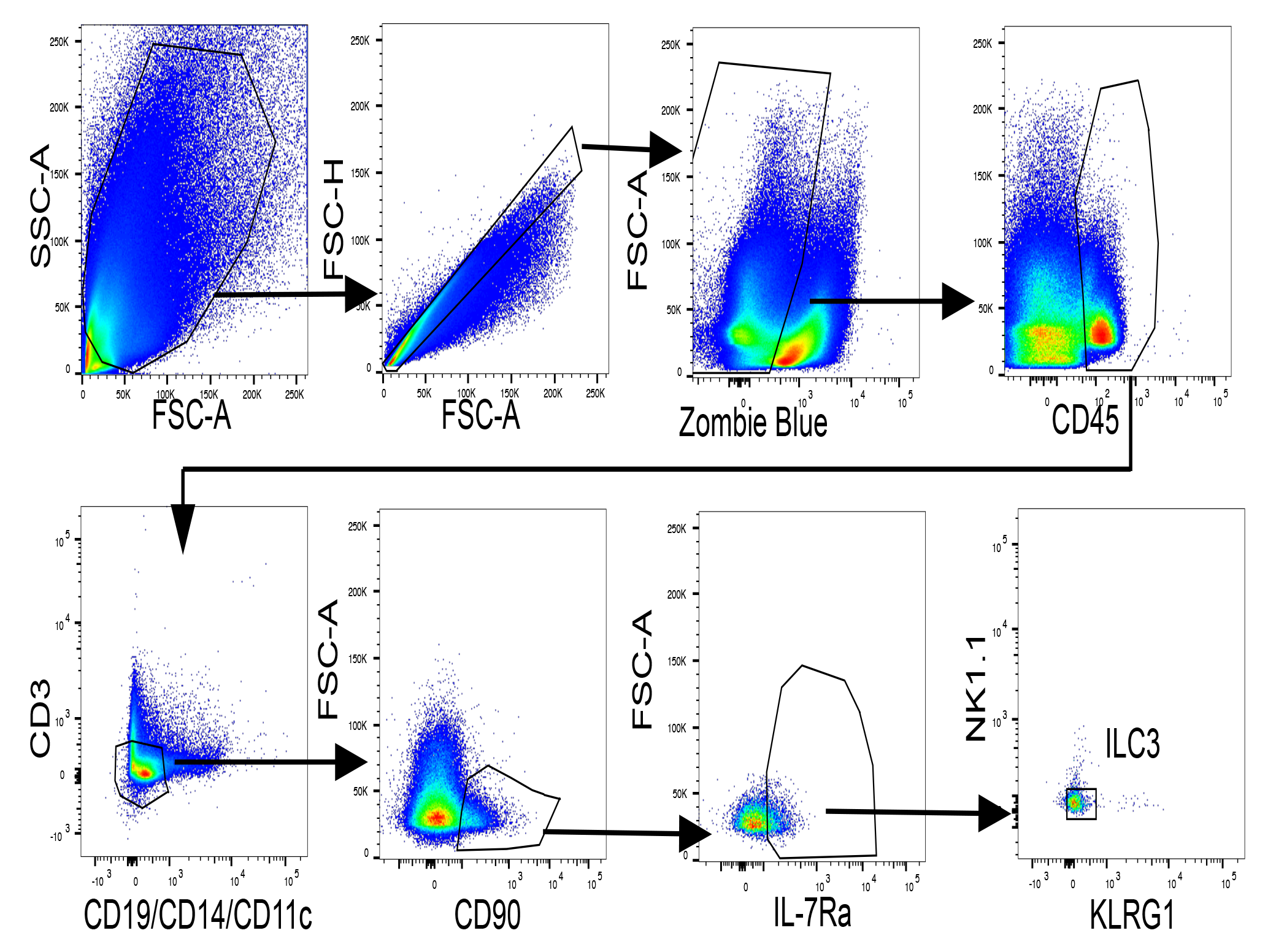
Schematic for determination of ILC3s within the lamina propria. iNLRC4*-Pou2f3-Cre^ERT2^* and *Nlrc4*^–/–^ littermate controls were retro-orbitally injected with two Flatox dosages 48 hours apart and small intestines were harvested 24 hours after last Flatox injection (LFn-FlaA (0.4ug/g) and PA (0.8ug/g)). Isolated lamina propria cells were stained with flow panel described in methods to identify ILC3s. ILC3s were characterized as CD45^+^, Lin^–^ (CD3, CD19^-^, CD14^-^, CD11c^-^), CD90^+^, IL7Ra^+^, NK1.1^-^ and KLRG1^-^.

**Supplemental Fig. 2.**
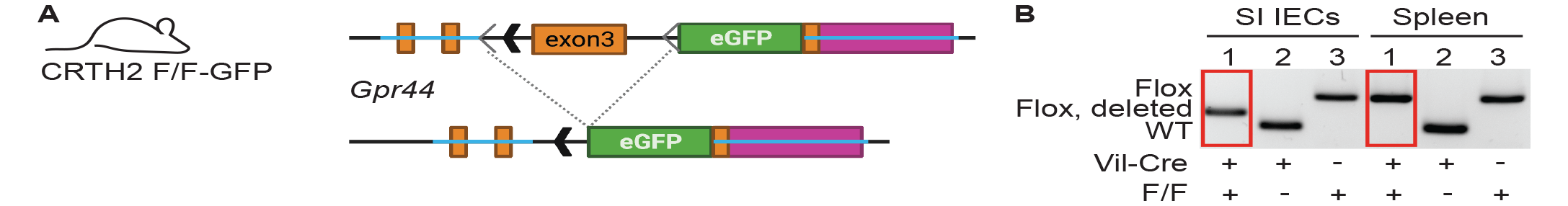
Validation of CRTH2^fl/fl^-GFP mice. (A) Schematic of mouse design. LoxP sites were inserted into the Gpr44 exon 3, followed by the sequence encoding eGFP. The presence of Cre recombinase excises the targeted segment of Gpr44 exon 3, resulting in deletion and allowing for transcription of eGFP from the endogenous Gpr44 promoter. (B) Genotyping PCR showing cell lineage-specific deletion of the targeted Gpr44 exon 3 sequence. Recombination occurs only in the presence of Cre in a cell lineage-specific manner in small intestine intestinal epithelial cells (IECs) but not splenocytes in Vil1-Cre CRTH2 F/F-GPF mice. WT allele = 239 bp; Flox allele = 348 bp; Flox allele after recombination = 279 bp.

**Supplemental Fig. 3.**
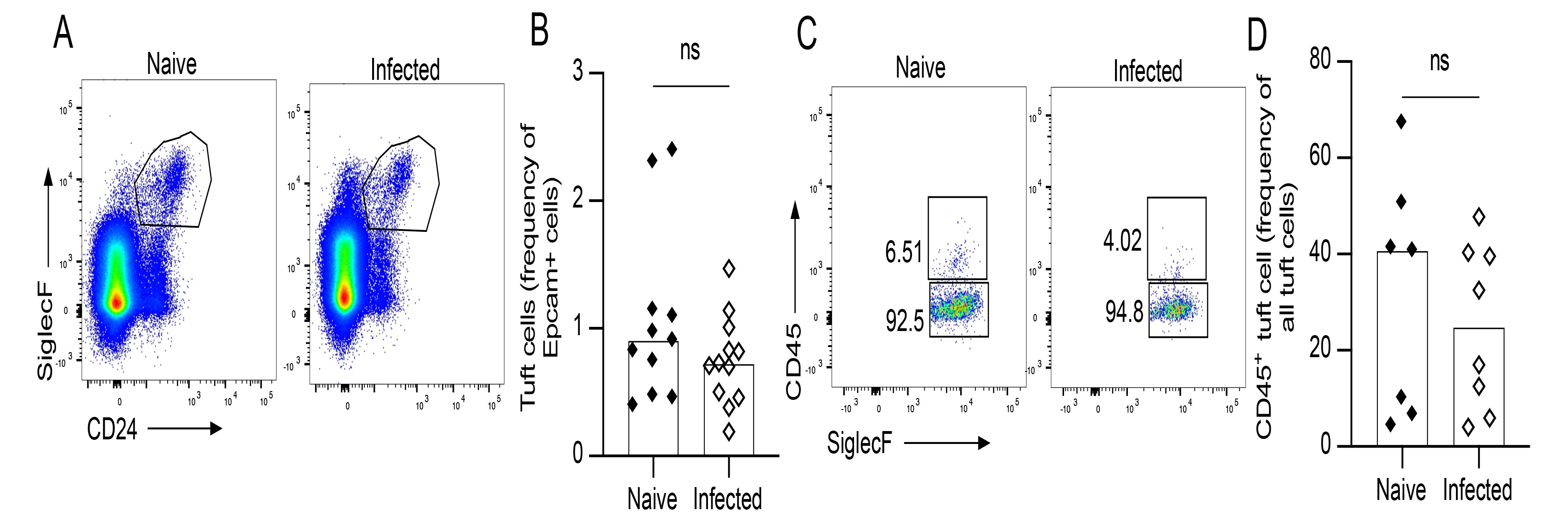
Tuft cell numbers are unchanged after *Shigella flexneri* infection. *Nlrc4*^+/–^ mice were pre-treated with oral gavage of streptomycin sulfate then infected with 5×10^6^ CFUs of *Shigella flexneri* by oral gavage. Two days post infection, the small intestine was harvested, and epithelial cells were isolated to be analyzed by flow cytometry. (A) Representative flow plots of total tuft cells from naïve and infected animals which is enumerated in (B). (C) Representative flow plots of CD45^+^ and CD45^-^ tuft cells, numbers are enumerated in (D). Graphs depict pooled data from three independent experiments, bars depict median. Significance was determined by Mann-Whitney U test.

